## Supplemental Figure 4 for "Regulatory context drives conservation of glycine riboswitch aptamers"

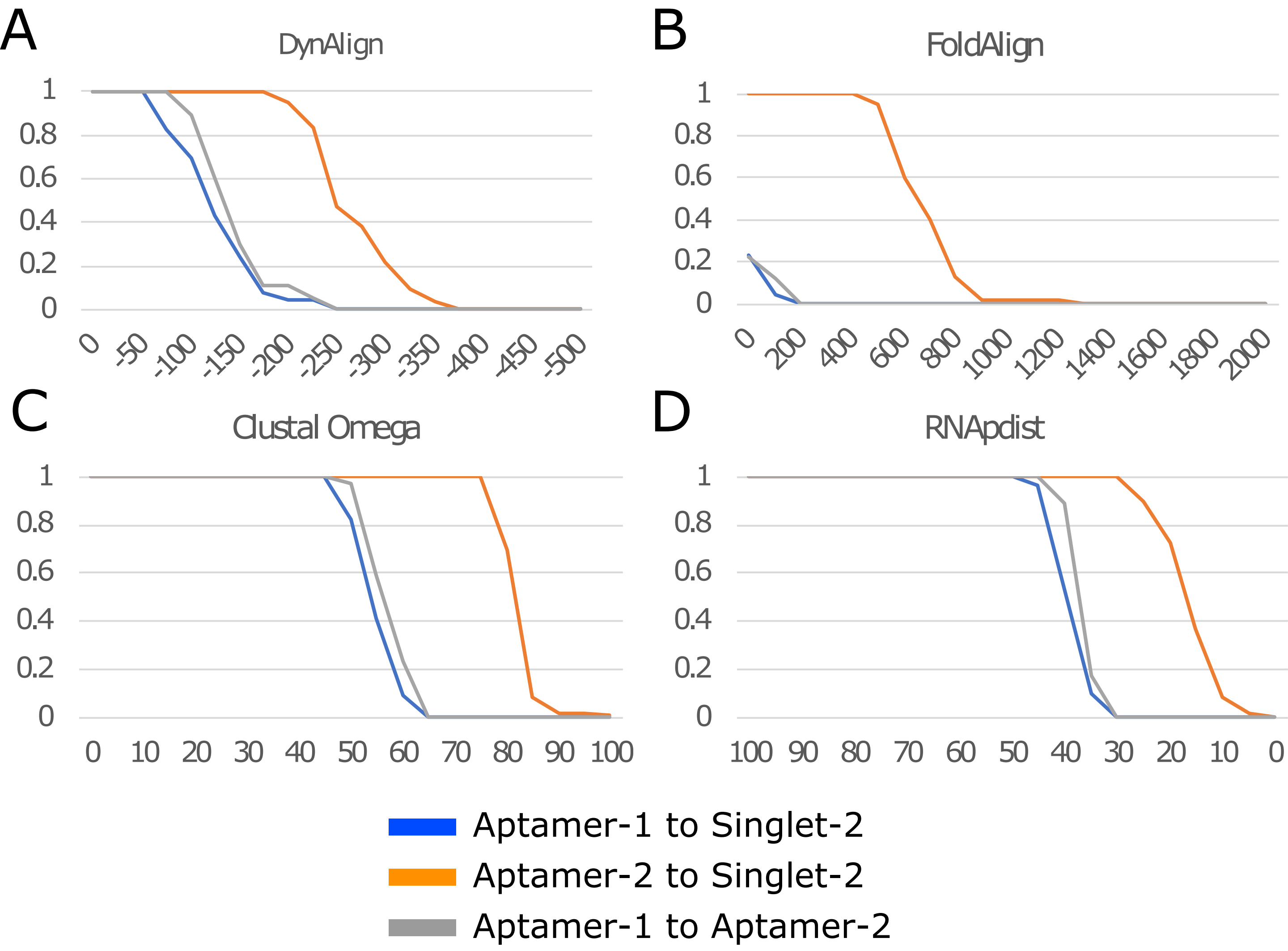


Supplementary Figure S4: Clustering of Bacilli aptamer-2 and singleton type-2 aptamer subset using Dynalign, FoldAlign, Clustal Omega, and RNApdist.
A) Dynalign inter-edge density across a range of -500 to 0 (x-axis reversed to display decreasing density).
B) FoldAlign inter-edge density across a range of 0 to 2000.
C) Clustal Omega inter-edge density across a range of 0 to 100.
D) RNApdist inter-edge density across a range of 0 to 100 (x-axis reversed to display decreasing density).
