## Supplemental Figure 7 for "Regulatory context drives conservation of glycine riboswitch aptamers"

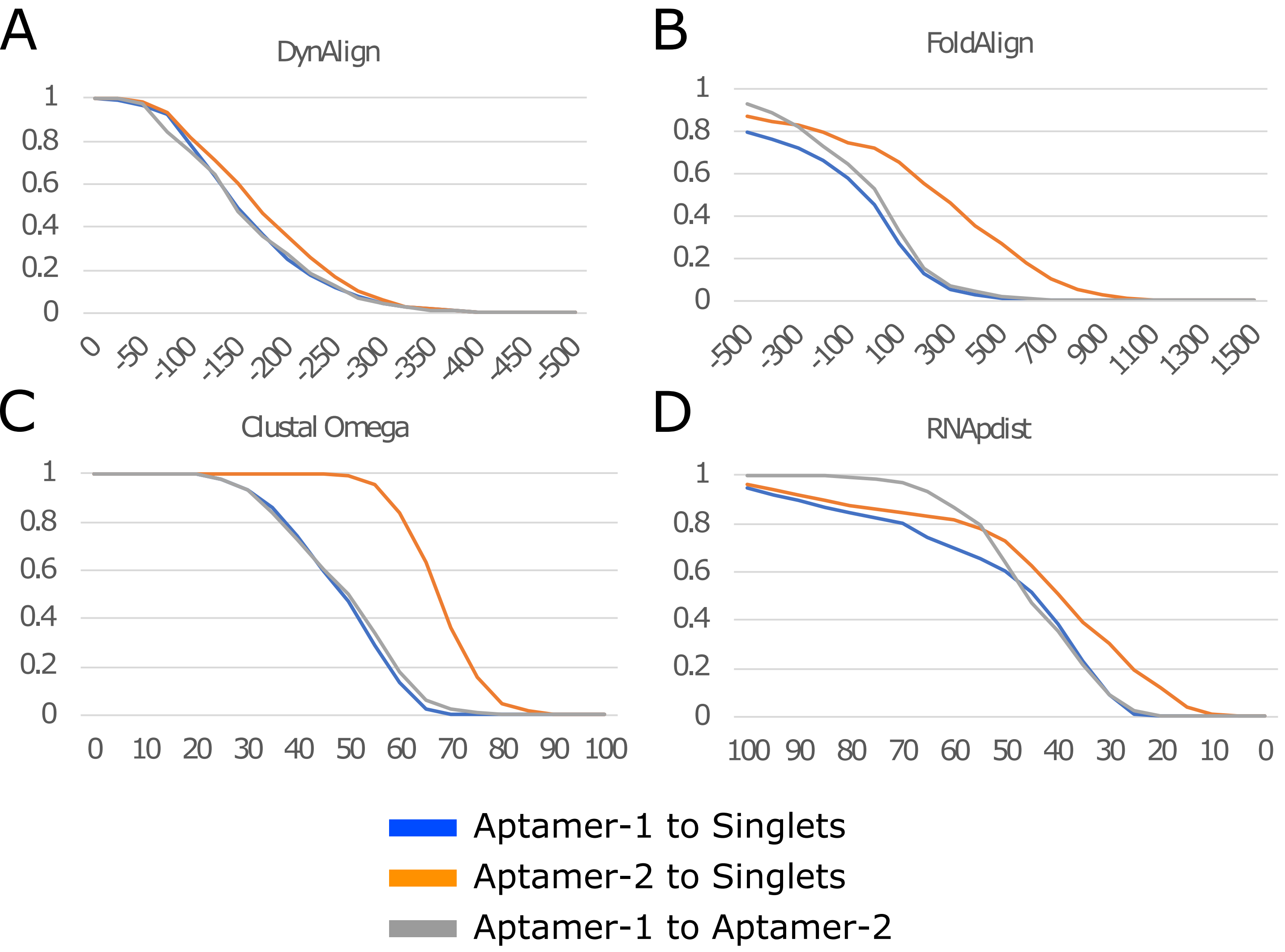


Supplementary Figure S7: Clustering of random riboswitch aptamers regulating TP using Dynalign, FoldAlign, Clustal Omega, and RNApdist.
A) Dynalign inter-edge density across a range of -500 to 0 (x-axis reversed to display decreasing density).
B) FoldAlign inter-edge density across a range of -500 to 1500.
C) Clustal Omega inter-edge density across a range of 0 to 100.
D) RNApdist inter-edge density across a range of 0 to 100 (x-axis reversed to display decreasing density).
